# Popeye Domain-Containing Protein 1 Scaffolds a Complex of Adenylyl Cyclase 9 and the Two-Pore-Domain Potassium Channel TREK-1 in Heart

**DOI:** 10.1101/2021.12.21.473719

**Authors:** Tanya A. Baldwin, Yong Li, Autumn Marsden, Roland F.R. Schindler, Musi Zhang, Anibal Garza Carbajal, Mia A. Garcia, Venugopal Reddy Venna, Thomas Brand, Carmen W. Dessauer

**Author notes:** Corresponding author information: Dr. Carmen W. Dessauer, Department of Integrative Biology and Pharmacology, McGovern Medical School, University of Texas Health Science Center, Houston, TX 77030. These authors contributed equally to this work (Tanya A. Baldwin, Yong Li).

## Abstract

The establishment of macromolecular complexes by scaffolding proteins such as A-kinase anchoring proteins is key to the local production of cAMP by anchored adenylyl cyclase (AC) and the subsequent cAMP signaling necessary for many cardiac functions. We have identified herein a novel AC scaffold, the Popeye domain-containing (POPDC) protein. Unlike other AC scaffolding proteins, POPDC1 binds cAMP with high affinity. The POPDC family of proteins are important for cardiac pacemaking and conduction, due in part to their cAMP-dependent binding and regulation of TREK-1 potassium channels. TREK-1 binds the AC9:POPDC1 complex and co-purifies in a POPDC1-dependent manner with AC9-associated activity in heart. Although the interaction of AC9 and POPDC1 is cAMP independent, TREK-1 association with AC9 and POPDC1 is reduced in an isoproterenol-dependent manner, requiring an intact cAMP binding Popeye domain and AC activity within the complex. We show that deletion of *Adcy9* (AC9) gives rise to bradycardia at rest and stress-induced heart rate variability. The phenotype for deletion of *Adcy9* is milder than previously observed upon loss of *Popdc1*, but similar to loss of *Kcnk2* (TREK-1). Thus, POPDC1 represents a novel scaffolding protein for AC9 to regulate heart rate control.

**ONE-SENTENCE SUMMARY:** Adenylyl cyclase type 9 binds in an isoproterenol-dependent manner to the POPDC1:TREK-1 complex regulating heart rate.

## INTRODUCTION

Cyclic AMP mediates the sympathetic regulation of many physiological functions in the heart, including contractility, relaxation, heart rate (HR), conduction velocity, and stress responses (*1*). The robust and specific control of so many activities by cAMP within a single cardiomyocyte and/or nodal cell is due not only to the diversity of adenylyl cyclase (AC) isoforms and cAMP effector molecules, but also to the spatial control of cAMP signaling by the generation of macromolecular complexes (*1-3*). These complexes are organized by a family of A-kinase anchoring proteins (AKAPs) that anchor protein kinase A (PKA) in close proximity to its targets for regulation by phosphorylation (*4*). Many of these complexes also contain AC and phosphodiesterase, providing a framework for integration and modulation of local cAMP signaling within distinct AC-PKA protein complexes or nanodomains (*5, 6*). There are 9 transmembrane AC isoforms that display diverse patterns of regulation, yet only a subset of AC isoforms bind to an individual AKAP (*7, 8*). For example, coupling AC5 with a downstream effector of PKA (i.e. TRPV1) on AKAP79 can sensitize this anchored effector to local cAMP production, reducing the IC_50_ for agonists by 100-fold (*9-11*). Scaffolding of AC9 to AKAP9 (a.k.a. Yotiao) promotes the PKA-dependent phosphorylation of KCNQ1 and is required for regulation of the I_Ks_ current in cardiomyocytes (*10, 11*).

We show herein that the <u>Pop</u>eye <u>d</u>omain <u>c</u>ontaining (POPDC) protein represents a novel scaffolding protein for AC isoforms. POPDC isoforms 1-3 were named for their conserved and abundant expression in skeletal and cardiac muscle and are important for cardiac pacemaking and conduction (*12*). POPDC proteins contain a short amino terminal extracellular domain, which is heavily glycosylated, three transmembrane domains and a cytosolic Popeye domain that displays structural similarity to the regulatory subunit of PKA and binds cAMP with high (∼120 nM) affinity (*13*). Loss of either *Popdc1* (a.k.a. *Bves*) or *Popdc2* in mice or zebrafish results in arrhythmic and bradycardic phenotypes with high HR variability (*13-15*), while human mutations of POPDC family members cause limb-girdle muscular dystrophy (LGMD), cardiac arrhythmia, familial atrioventricular (AV) block, and are implicated in long-QT syndrome and heart failure (*15-23*). POPDC1 and POPDC2 bind the two-pore-domain potassium channel TREK-1 (KCNK2 or K2P2.1) to enhance channel density at the plasma membrane and increase K^+^ currents (*13, 15, 16*). Upon cAMP binding to POPDC proteins, the POPDC:TREK-1 complex is dissociated (*13, 16*); loss of this regulation is proposed in part to give rise to HR variability upon deletion of *Popdc1* or *-2*.

AC9 also regulates HR. Deletion of AC9 gives rise to a bradycardia at rest and HR variability during recovery from stress (*24*), albeit with a milder phenotype than loss of POPDC1. We show that AC9 binds all three isoforms of POPDC, interacting with both the transmembrane regions and the cytosolic Popeye domain of POPDC1. TREK-1 co-localizes and associates with the AC9:POPDC complex, while the deletion of AC9 or POPDC1 reduces TREK-1 associated Gαs-stimulated AC activity in heart. TREK-1 also interacts with the calcium calmodulin-stimulated ACs (AC1 and AC8), however this interaction is independent of POPDC1. Binding of AC9 and POPDC is independent of cAMP production, while AC9 association with TREK-1 is reduced in an isoproterenol (ISO)-dependent manner, requiring an intact Popeye domain and local production of cAMP within the complex. POPDC1 therefore represents a novel scaffolding protein for AC9 to regulate downstream effectors for HR control.

## RESULTS

### Deletion of AC9 decreases HR at rest and increases HR variability after ISO injection

Previous studies using Doppler imaging of isoflurane-anesthetized mice revealed a mild bradycardia in *Adcy9*^-/-^ male and female mice from 1-7 months of age (*24*). To confirm effects of AC9 deletion on HR in conscious animals, we measured electrocardiograms (ECGs) by telemetry. HR was measured over a 24-hour period in which the animals were given free access to a running wheel. *Adcy9*^-/-^ mice displayed a daytime bradycardia during periods of rest (defined as no movement of the running wheel for >10 min), but not while actively moving (>5 min on running wheel) (Fig 1A). Similar trends were also observed at night.

**Fig. 1.**
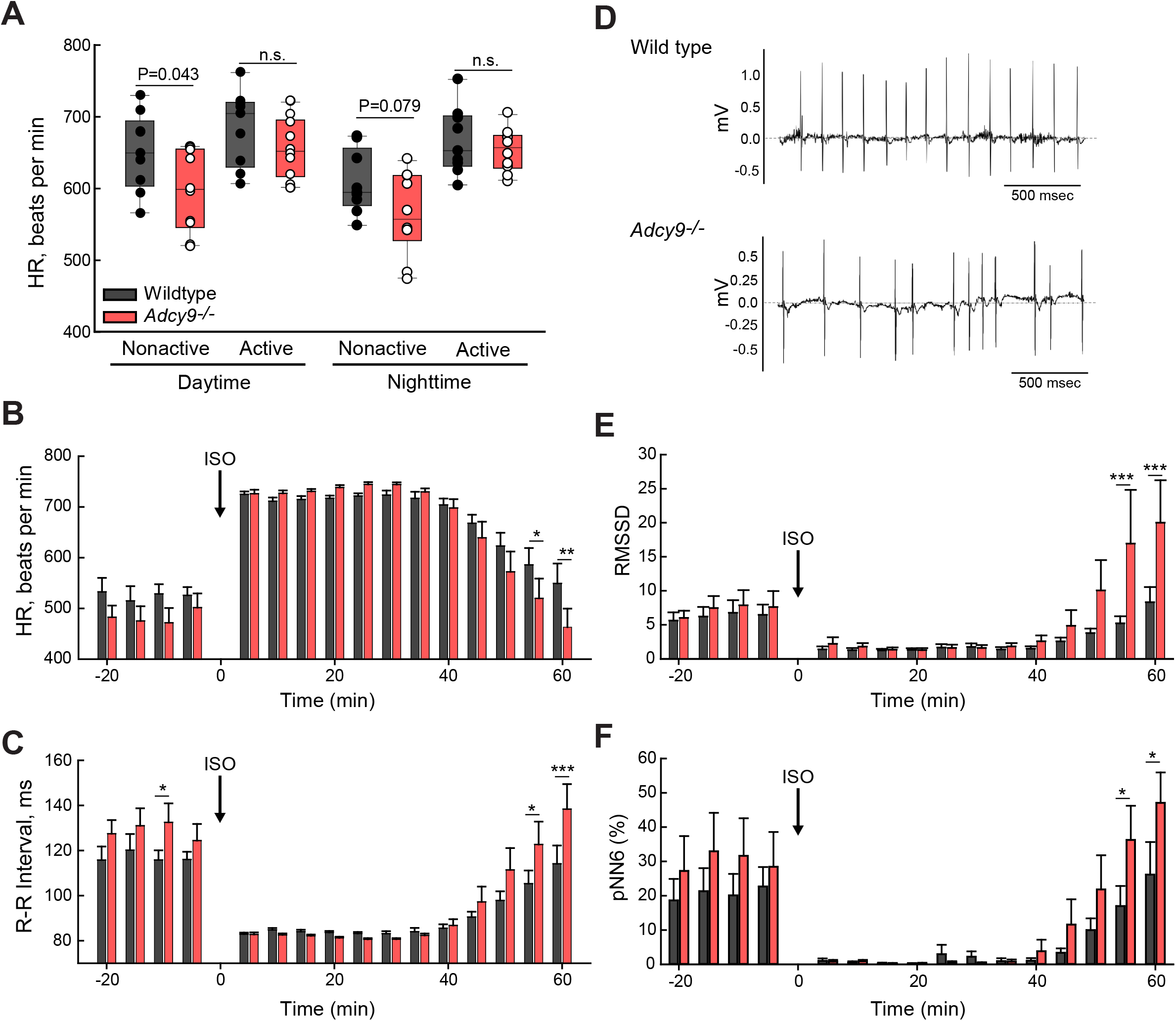
Deletion of *Adcy9* results in bradycardia at rest and HR variability during recovery period after ISO injection. (**A**) Heart rate (HR) of WT and *Adcy9*^-/-^ (AC9 KO) mice averaged over 5 min of non-active versus sustained active periods on home cage running wheels. Student’s t-test P values for indicated comparisons are shown (n=9 WT and n=10 *Adcy9*^-/-^ mice (4 females, 6 males). (**B**,**C**) HR and R-R interval of WT (black bars) and *Adcy9*^-/-^ mice (red bars) before and after ISO (1 μg/g) injection; mean of 5 min intervals +/- SE are shown. (**D**) Representative ECG recordings of 7 mo WT and *Adcy9*^-/-^ mice, ∼55 min post ISO-injection. (**E, F**) HR variability was calculated before and after ISO injection using the root mean square of successive RR interval differences (RMSSD; panel (**E**)) and percentage of sequential R-R intervals differing by >6 ms (pNN6; panel (**F**)) methods. Two-way ANOVA Repeated measures analysis (time and genotype) was performed with Holm-Sidak method used for pairwise comparisons for HR, RR, and HR variability with ISO; n=9 WT and n=10 *Adcy9*^-/-^ mice; *P<0.05; **P<0.01; ***P<0.001).

Mice were subjected to behavioral tests post implantation of telemetry devices to rule out effects on general locomotor activity levels and exploratory drive (running wheels and open field activity box) or anxiety (elevated plus maze and forced swim test) that may affect overall HR. Overall, no significant differences were found in the maximal velocities or total distance traveled on the running wheel or during the 25 min open field test (Table 1). *Adcy9*^-/-^ also showed no exploratory differences in the open field test and displayed similar levels of anxiety and depression-like behavior as demonstrated in the elevated plus maze and forced swim tests (Table 1).

**Table 1.**
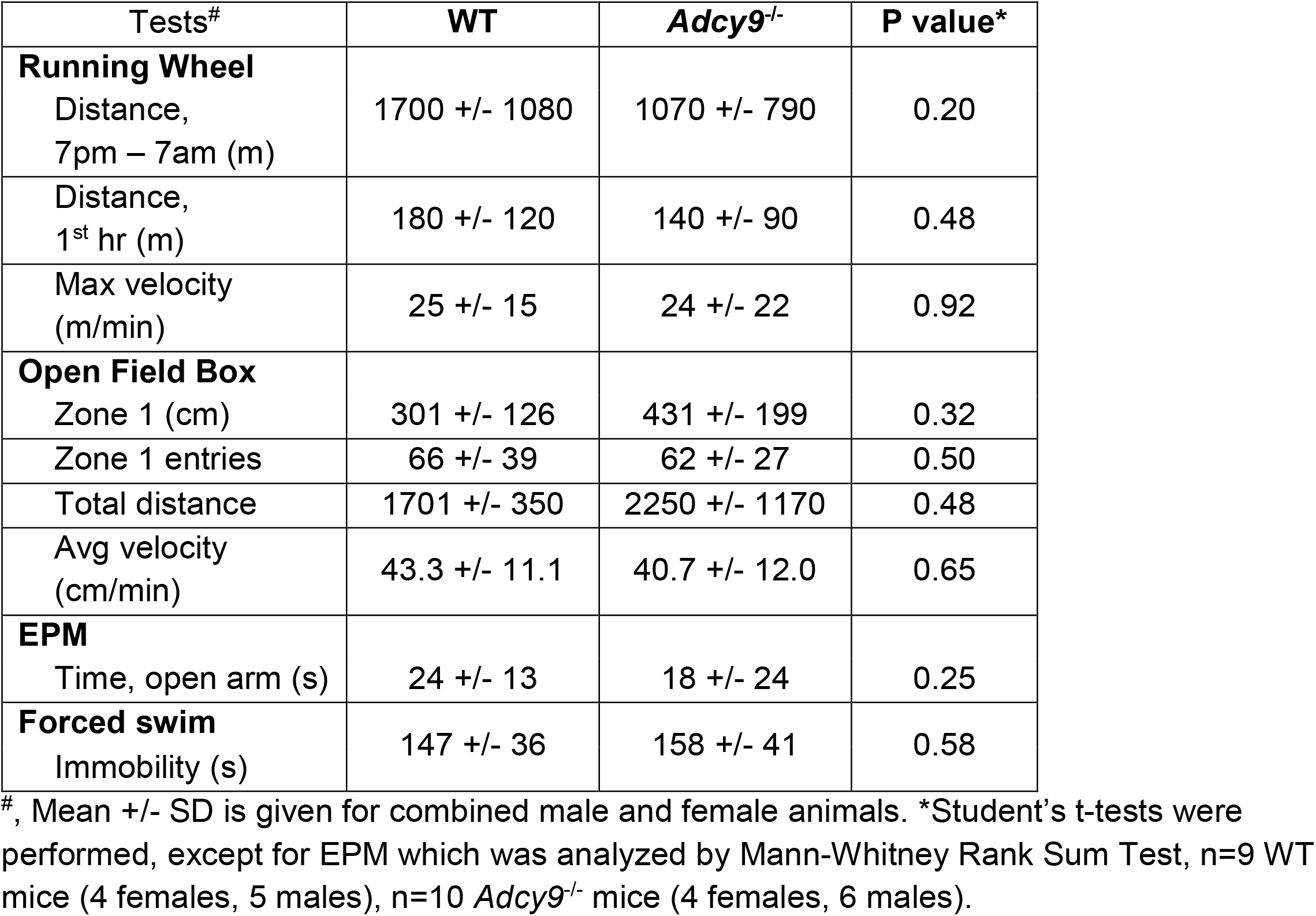
Behavioral assays for WT and *Adcy9*^-/-^ mice.

To induce an increase in HR in a controlled and timed manner, we administered ISO (1 μg/g) by a single intraperitoneal injection and recorded ECGs. Both genotypes and sexes showed a substantial increase in HR upon ISO administration (39 +/- 10% and 48 +/- 23% for WT and KO, respectively; Fig 1B), reaching a similar maximal HR. However, *Adcy9*^-/-^ displayed a more rapid recovery to baseline with lower HR and higher R-R intervals at 55 and 60 min post injection (Fig 1B, C). A further examination of ECG traces revealed a variability in beat-to-beat timing in *Adcy9*^-/-^ mice at these time points (Fig 1D). HR variability was quantified by two distinct methods (RMSSD and pNN6); both methods showed statistically significant HR variability in the recovery phase after ISO injection (Fig 1E, F).

### POPDC proteins bind AC9

Despite the effects on HR and HR variability during recovery from beta-adrenergic stimulation, it was not immediately clear why deletion of AC9 would give rise to this phenotype. AC9 is present in sinoatrial (SA) node (*24*), but the Ca^2+^/calmodulin-stimulated ACs are generally thought to be responsible for pacemaker activity within the SA node (*25-28*). Moreover, numerous ion channels associate with the AKAP scaffold AKAP79/150 to regulate HR, but AC9 contributes little to no activity within this complex in heart (*24*). Thus we searched the literature for proteins that were regulated by cAMP and when deleted, displayed HR variability. Two proteins of interest included POPDC1 and TREK-1; upon deletion, both proteins show a sinoatrial phenotype and HR variability (*13, 29*). Additionally, a high-throughput affinity purification-mass spectrometry screen identified AC9 and AC3 with POPDC2 as bait, however these potential interactions were never biochemically verified (*30*). POPDC proteins represent a family of cAMP effectors that bind to the two-pore potassium channel TREK-1 in a cAMP-dependent manner (*18, 31*).

Unfortunately, AC9 and POPDC1 suffer from a lack of antibodies sufficient for immunoprecipitation, with poor detection of endogenous proteins by western blotting. Therefore, we first tested for potential interactions of AC9 and POPDC using expression of tagged proteins in cellular assays of HEK293 cells and COS-7 cells (Fig 2). Proximity ligation assay (PLA) can amplify detection of protein-protein interactions that occur within a range of <60 nm (*32*). YFP-tagged AC9 and POPDC-Myc tagged proteins were expressed in HEK293 cells; interactions between AC9 and Gβγ (as detected with antibodies against Gβ) were used as a positive control (*24*). PLA puncta (red), as quantitated by high content microscopy, are readily detected with AC9 and POPDC isoforms 1 and 2, suggesting close proximity and possible complex formation (Fig 2B, C). A complementary cellular interaction technique is Bimolecular Fluorescence Complementation (BiFC). Here, potential interacting pairs are tagged with either the N-terminal half of Venus (VN) or C-terminal half of Venus (VC). Since HEK293 cells express endogenous POPDC1 and POPDC2, we performed BiFC experiments in COS-7 cells that lack detectable endogenous POPDC proteins. A strong BiFC signal is detected between AC9 and POPDC 1 and 3, and to a lesser extent with POPDC2 (Fig 2D), with AC9:POPDC1 BiFC signal localizing to PM in HEK293 cells (Fig 2E).

**Fig. 2.**
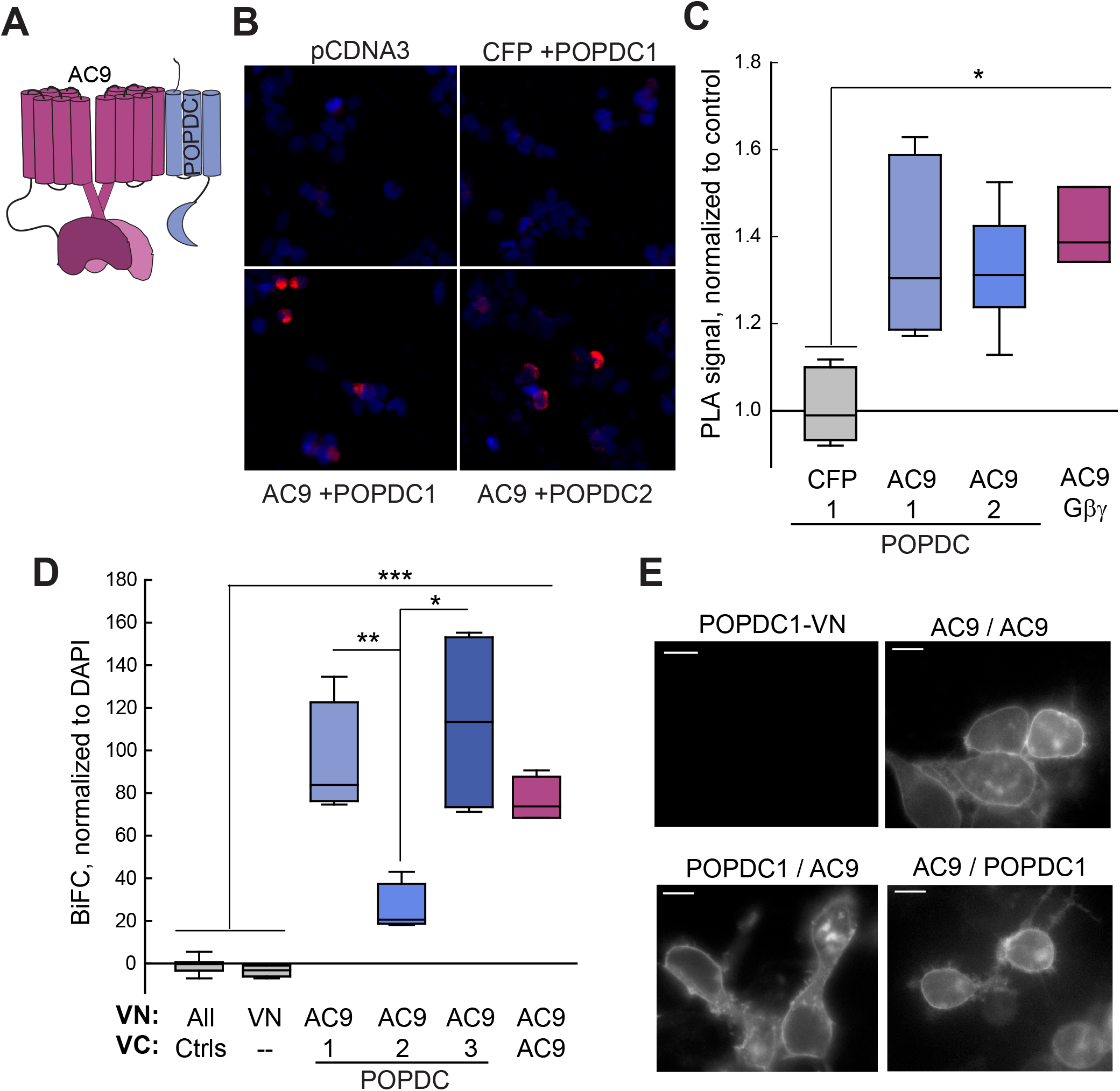
Cellular interaction of POPDC isoforms with AC9. (**A**) Cartoon of AC9-POPDC complex. (**B**,**C**) Proximity ligation assay was performed with HEK293 cells expressing pCDNA3 (background control), POPDC1-Myc alone, and YFP-AC9 in the presence of either POPDC1-Myc or POPDC2-Myc. The interaction between AC9 and Gβγ served as a positive control (*69*). (**B**) Images of PLA signal (red) and DAPI (blue). (**C**) Mean cellular fluorescence intensity was quantified by high content microscopy and shown as Box and Whisker plots. Kruskal-Wallis One Way ANOVA analysis was performed (n=7 experiments, P=0.003 between groups) with multiple comparisons by Bonferroni t-test (*P<0.05) (**D**) Quantification of COS7 cells expressing BiFC constructs for AC9, POPDC 1-3. The expressed proteins tagged with VN (top line) and VC (bottom line) are shown. Kruskal-Wallis One Way ANOVA analysis was performed (n=4 experiments, ***P<0.001 between groups) with multiple comparisons to control by Dunn’s method (**P<0.01, *P<0.05) (**E**) Representative live-cell images of indicated BiFC combinations in HEK293 cells (n>20 cells; scale bar is 10 um). Quantification of POPDC1-VN:AC9-VC is shown in Fig 3 and 5.

To test whether AC and POPDC interactions could also be detected by co-immunoprecipitation (co-IP), Flag-tagged AC9 was co-expressed with or without Myc-tagged POPDC1 or POPDC2 in HEK293 cells. Complexes were isolated with anti-MYC antibodies and associated Gαs-stimulated AC activity was measured (IP-AC assay; Fig 3A). From the IP-AC assay it appeared that AC9 interacted with POPDC1 but not POPDC2. However, when the co-immunoprecipitations were evaluated via western blot, AC9 co-precipitates with both POPDC1 and POPDC2 (Fig 3B). Thus, POPDC1 pulls down AC9 and maintains associated AC activity, while AC9 complexed with POPDC2 does not respond to Gαs stimulation. POPDC1/2 interactions with AC9 were confirmed by performing co-immunoprecipitations in reverse by pulling down the complex with the associated Flag-tag on AC9 (Fig 3C). POPDC1 and POPDC2 were both detected in pull-downs of Flag-AC9.

**Fig. 3.**
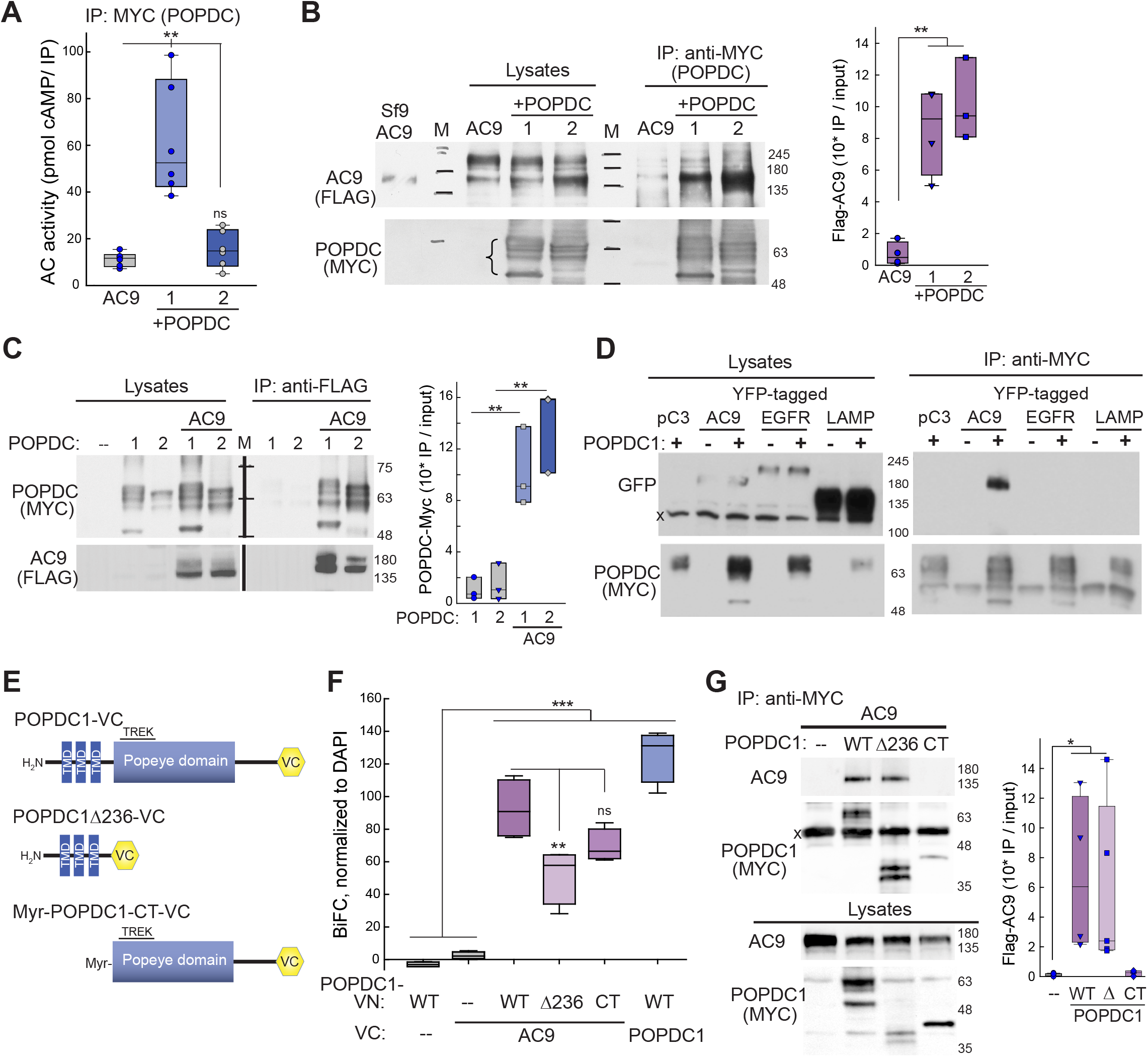
AC9 interacts with the POPDC1 transmembrane and Popeye domains. (**A**) Cell lysates from HEK293 cells expressing Flag-AC9 in the presence or absence of Myc-tagged POPDC1 or −2 were subjected to co-immunoprecipitation (Co-IP) with anti-MYC and assayed for AC activity with 300 nM Gαs-GTPγS. Kruskal-Wallis One Way ANOVA analysis on Ranks was performed (n=6 experiments, P=0.003 between all groups) with multiple comparisons to AC9 control by Dunn’s method (**P<0.01). (**B**) A portion of the lysates and Co-IP from (**A**) were subjected to western blot (WB) analysis with anti-MYC (POPDC) and anti-FLAG (AC9). Membranes isolated from Sf9 cells expressing Flag-AC9 served as a positive WB control. Note, POPDC protein runs as multiple bands (48-65 kDa) with altered sizes/patterns in different tissues due to changes in glycosylation patterns (*12*). Molecular weight markers are denoted as M. Quantitation of Flag-AC9 WB by One Way ANOVA, n=3-4 experiments, with comparisons by Tukey test, **P<0.01). (**C**) HEK293 cells expressing Flag-AC9 +/- POPDC1-Myc or POPDC2-Myc were subjected to Co-IP with anti-FLAG and subjected to WB analysis with anti-MYC and anti-FLAG. Quantitation of anti-MYC WB by One Way ANOVA, n=3 experiments, with comparisons by Tukey test, **P<0.01). (**D**) POPDC1 does not interact with control TM proteins. HEK293 cells expressing POPDC1-Myc +/- GFP-tagged AC9, EGFR, or LAMP1 were subjected to Co-IP with anti-MYC. Western blotting of lysates and Co-IPs for GFP (top) and Myc (bottom) are shown (n=3 experiments). (**E**) Schematic of POPDC1 truncations. (**F**) BiFC of AC9 and indicated POPDC1 truncations in COS-7 cells. Kruskal-Wallis One Way ANOVA analysis was performed (n=4 experiments, ***P<0.001 between groups) with multiple comparisons by Tukey test (**P=0.008). (**G**) Co-IP with anti-MYC in COS-7 cells expressing Flag-AC9 and indicated Myc-tagged POPDC1 truncations. Western blotting with anti-AC9 and anti-MYC (POPDC1) of Co-IP and lysates is shown. Kruskal-Wallis One Way ANOVA analysis was performed (n=3-5 experiments, P=0.003), with multiple comparisons by Dunn’s method (*P<0.05).

A number of POPDC interacting proteins have been identified to date, many of which are transmembrane proteins (reviewed in (*33*)). To verify the specificity of our immunoprecipitation conditions, we confirmed the association of POPDC1 with AC9, but not the transmembrane epidermal growth factor receptor (EGFR) or lysosomal associated membrane protein 1 (LAMP1) (Fig 3D). To begin to narrow down the site of AC:POPDC interaction, the transmembrane domain (POPDC1Δ236) or the PM-targeted cytosolic Popeye domain (Myr-POPDC1-CT) were tested for interactions with AC9 (Fig 3E-G). Both the transmembrane and cytosolic domain of POPDC1 interacted with AC9 by BiFC, although deletion of the cytosolic domain gave rise to a reduced BiFC signal compared to the full-length protein. This may be due to alterations in the distance between VN and VC proteins with the truncated POPDC1, as POPDC1Δ236 interacted with AC9 by co-immunoprecipitation, similar to full-length POPDC1. Interactions between AC9 and the cytosolic domain of POPDC1 were not observed by pull-down, suggesting that although both domains participate in interactions with AC9, the stronger interaction likely occurs within the transmembrane spanning regions.

### POPDC1 mediates interaction of TREK-1 with AC9

POPDC expression enhances TREK-1 K^+^ currents, while loss of POPDC-regulated TREK-1 currents is considered to be at least one underlying cause of the stress-induced bradycardia observed in *Popdc1*^-/-^ or *Popdc2*^-/-^ mice (*13*). TREK-1 co-localized in HEK293 cells with the BiFC signal from the AC9 homodimer (Fig 4A, top) and the BiFC complex of POPDC1:AC9 (Fig 4A, bottom), suggesting that all three proteins co-localize on the plasma membrane. Cellular interactions can be observed by fluorescence lifetime microscopy (FLIM) of Cerulean-tagged proteins to quantitatively measure FRET efficiency with YFP-tagged partners (Fig 4B). Using this method, interactions were observed between TREK-1 and POPDC1 (18 +/- 2% FRET efficiency) and POPDC1 and AC9 (6.2 +/- 0.4% FRET). The low FRET efficiency is likely due to the distance constraints of FRET, given that all three proteins (TREK-1, POPDC1 and AC9) form dimers. Despite the low FRET efficiency, immunoprecipitation of POPDC1 pulled down both AC9 and TREK-1 when co-expressed in HEK293 cells (Fig 4C). Similarly, immunoprecipitation of TREK-1 brought down AC9 and endogenous or overexpressed POPDC1 (Fig 4D). TREK-1 interacts with AKAP79 (*34*); however, overexpression of AKAP79 did not enhance AC9 pull-down with TREK-1, nor was AKAP79 detected in immunoprecipitates of TREK-1 (Fig 4D).

**Fig. 4.**
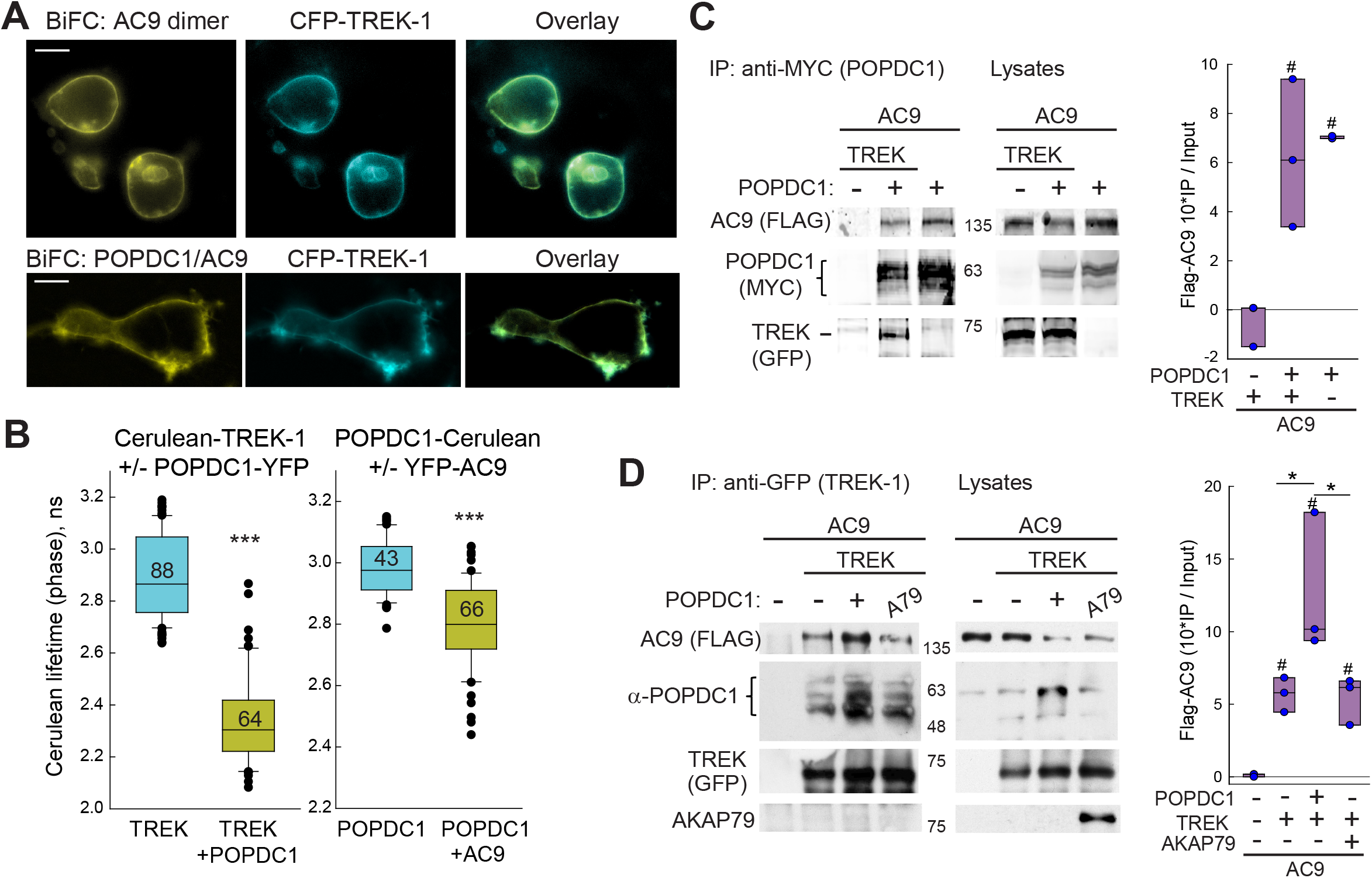
AC9 is part of a POPDC1-TREK-1 complex. (**A**) Co-localization of CFP-TREK-1 with the BiFC signal from AC9-VN:AC9-VC homodimer (top) and POPDC1-VN:AC9-VC complex (bottom) in HEK293 cells. Scale bar is 10 um. (**B**) The lifetime distribution of Cerulean-tagged proteins +/- the indicated YFP-tagged proteins expressed in HEK293 cells are displayed as box and whisker plots with all outliers shown. Mann-Whitney Rank Sum Test was performed (***P<0.001; n=cell number indicated on each bar). (**C**) Co-IP of Myc-tagged POPDC1 (anti-MYC) pulls down both Flag-AC9 and GFP-tagged TREK-1 (n=3 experiments). (**D**) Co-IP of TREK-1 (anti-GFP) pulls down Flag-AC9 and endogenous POPDC1. AKAP79 is not pulled down in complex (n=3 experiments). Quantification of Flag-AC9 WB (IP/total expression in lysate) is shown to the right of (**C**) and (**D**).

### AC9 interacts with TREK-1 in an ISO-dependent manner that requires POPDC1

Binding of POPDC proteins to TREK-1 promotes the membrane trafficking of the two-pore potassium channel and thus enhances K^+^ currents, while cAMP binding to POPDC proteins decreases TREK-1:POPDC interactions (*13*). Therefore, we examined the effect of ISO-stimulation on complex formation between AC9, POPDC1, and TREK-1 using BiFC. Interactions between TREK-1 and AC9 or TREK-1 and POPDC were examined by BIFC, although the signals were generally lower for these interactions as compared to AC9:POPDC1 or dimer formation by TREK-1, AC9, or POPDC1 (Fig 5A). Upon ISO treatment (10 min, 10 μM), the overall BiFC signal for TREK-1 and AC9 or TREK-1 and POPDC1 was decreased approximately 50-65%, independent of which protein was tagged with the amino or carboxy-terminal half of Venus (Fig 5B). The IC_50_ for the ISO-induced decrease in TREK-1:AC9 BiFC signal was 160 +/- 40 nM (Fig 5C). ISO regulation of TREK-1 required both an active AC9 enzyme and endogenous expression of POPDC1 (Fig 5C-D). TREK-1 interactions with a catalytically dead AC9 were not sensitive to ISO, suggesting that local cAMP production was important (Fig 5C). Significant endogenous expression of POPDC1 was also required for ISO regulation of TREK-1 interactions, as no significant TREK-1:AC9 BiFC signal was observed in COS-7 cells lacking detectable POPDC1 (Fig 5D). Finally, an intact Popeye domain was required, as overexpression of a truncated POPDC1 protein (POPDC1Δ172), which lacks cAMP binding but retains TREK-1 association (*15*), acted as a dominant negative to block the effects of ISO (Fig 5E). Expression of WT POPDC1 had no effect on TREK-1:AC9 BiFC signals. The interaction between AC9 and POPDC1 showed either no change or a 24 +/- 3% decrease with ISO, depending on which protein was tagged with VN and VC (Fig 5B). This dependence on VN/VC tagging may represent a change in orientation between the two proteins upon either Gαs interaction with AC9 or POPDC1’s binding of cAMP. Neither AC9 nor POPDC1 homodimers showed alterations in BiFC responses with ISO treatment (Fig 5B). This data support a model where local cAMP production by AC9 enhances POPDC1 binding of cAMP to drive the breakdown of the AC9:POPDC:TREK-1 assembly, releasing TREK-1 from the more stable AC9:POPDC1 complex (Fig 5F).

**Fig. 5.**
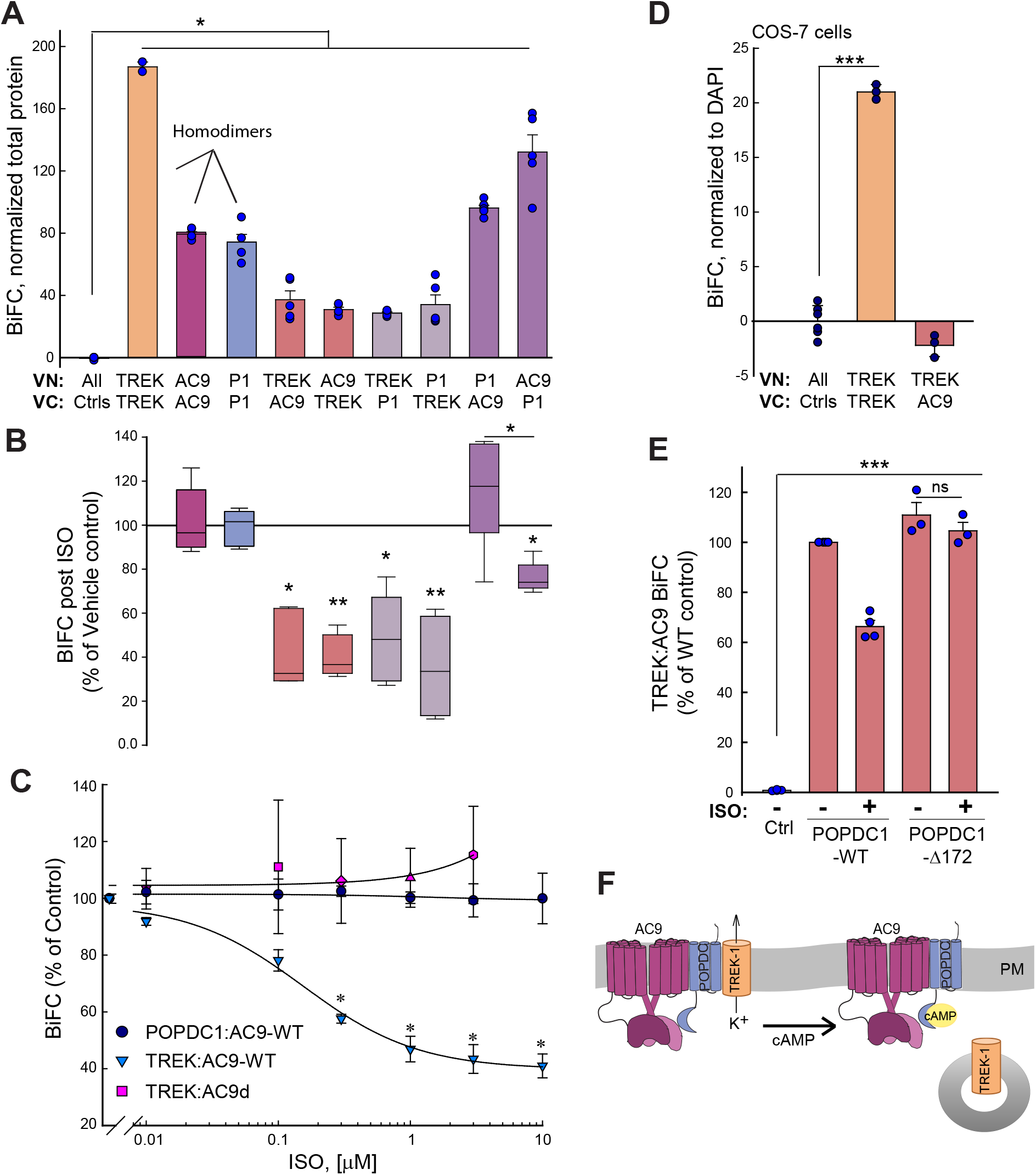
ISO-dependent loss of TREK-1 from AC9-POPDC1 complex requires an intact Popeye domain. (**A**) BiFC signal between the indicated VN- and VC-tagged proteins expressed in HEK293 cells (POPDC1, P1). Kruskal-Wallis One-Way ANOVA on Ranks (n=5 experiments, P<0.001 between groups) with multiple comparison to VN and VC controls by Student-Newman-Keuls tests (*P<0.05). (**B**) Percent decrease of BiFC signal in (**A**) with ISO treatment (10 μM, 10 min at 37°C). Box and whisker plots are shown. Paired t-test was performed for each condition (vehicle versus ISO) using raw data prior to calculation of percent of vehicle control (n=6; *P<0.05; **P<0.01). (**C**) ISO dose-response curves for BiFC interactions between POPDC1:AC9, TREK-1:AC9, and TREK-1 with a catalytically inactive AC9 (TREK-1:AC9d). Data were analyzed by Two-way ANOVA with multiple comparisons by Holm Sidak method (n=3-4; *P<0.05). (**D**) TREK-1:AC9 BiFC signal is absent in COS-7 cells lacking detectable POPDC1 (TREK-1:TREK-1 BiFC is shown as a positive control). One-way ANOVA with multiple comparisons by Holm Sidak method (n=3 experiments; ***P<0.001); (**E**) Overexpression of POPDC1 truncation of Popeye domain (POPDC1-Δ172), but not WT POPDC1, abolishes ISO reduction of TREK-1:AC9 BiFC signal. One-way ANOVA with multiple comparisons by Holm Sidak method (n=3-4 experiments; ***P<0.001 compared to control). (**F**) Model of cAMP effects on AC9-POPDC1-TREK-1 complex formation.

### Association of TREK-1 with AC9 activity in heart requires POPDC1 expression

To further validate the interaction between TREK-1 and AC9 in cardiac tissue, we measured the AC activity that was associated with immunoprecipitates of TREK-1 in hearts from WT or *Adcy9*^-/-^ mice. TREK-1 antibodies pulled down 8 +/- 2-fold more Gαs-stimulated AC activity than IgG controls in heart from WT mice, while AC activity associated with TREK-1 pulldown was significantly reduced, but not eliminated, in *Adcy9*^-/-^ mice (2.2 +/- 0.6-fold compared to IgG controls) (Fig 6A). Note that in WT and *Adcy9*^-/-^ mice, the total bulk AC activity and AKAP150-associated AC activity was unchanged (Fig 6A and (*24*)). POPDC1 was required for the association of Gαs-stimulated AC activity with TREK-1, as deletion of POPDC1 nearly eliminated all TREK-1 associated AC activity (Fig 6B). However, this was not due to alterations of total AC activity in *Popdc1*^-/-^ hearts (Fig 6C).

**Fig. 6.**
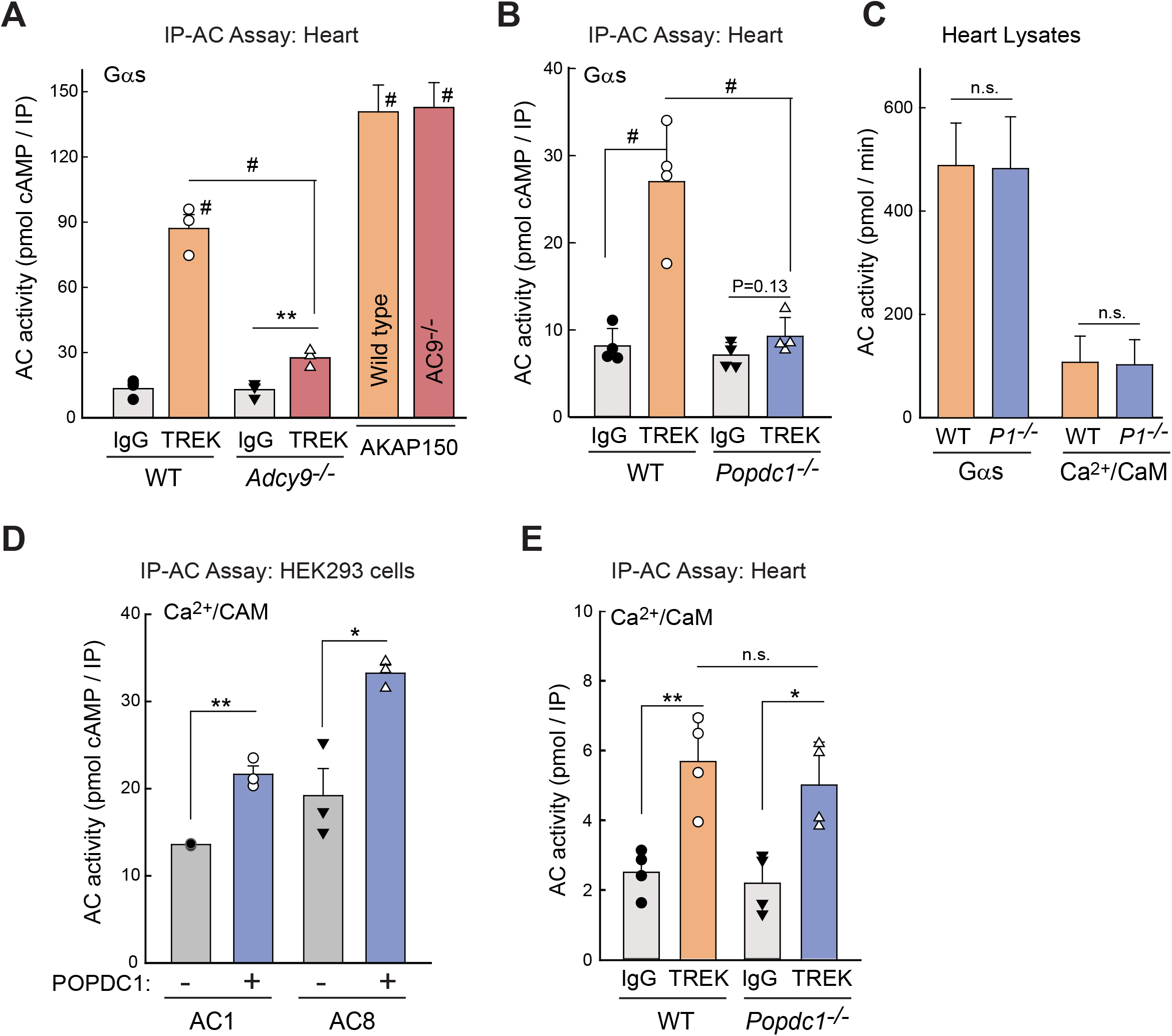
TREK-1 pulls-down AC9 activity in heart and interacts with Ca^2+^/calmodulin-stimulated AC1 and AC8. (**A**) IP-AC assay with IgG versus anti-TREK-1 in heart homogenates from WT and *Adcy9*^-/-^ mice; AC activity stimulated with 300 nM Gαs-GTPγS. IP-AC with anti-AKAP150 is shown as a positive control. Two-way ANOVA (overall P<0.001, n=3 mice per genotype) with multiple comparisons by Holm-Sidak test; ^#^P<0.001 for indicated comparison or to respective IgG controls. IgG versus anti-TREK-1 in *Adcy9*^-/-^ was analyzed by Student’s t-test (**P=0.009, n=3). (**B**) IP-AC assay with IgG versus anti-TREK in WT and *Popdc1*^-/-^ mouse heart homogenates; AC activity stimulated as in (A). Statistics as in A, n=4 mice per genotype, #P<0.001. Paired t-test within *Popdc1*^*-/*-^, P=0.123. (**C**) Total AC activity in heart homogenates from WT or *Popdc1*^*-/-*^ (*P1*^*-/-*^) mice stimulated with 300 nM Gαs-GTPγS or 100 μM calcium and 300 nM calmodulin. Analyzed by Student’s t-test (n=4, n.s.) (**D**) Cell lysates from HEK293 cells expressing AC1 or AC8 in the presence or absence of Myc-tagged POPDC1 were subjected to co-IP with anti-MYC and assayed for AC activity with 100 μM calcium and 300 nM calmodulin. Student’s t-test (n=3 experiments, **P=0.0011 and *P=0.012 for AC1 and AC8, respectively). **(E)** IP-AC assay with IgG versus anti-TREK-1 in WT and *Popdc1*^*-/-*^ mouse heart homogenates; AC activity stimulated 100 μM calcium and 300 nM calmodulin. Student’s t-test (*P=0.03; **P=0.005, n=4 mice per genotype).

### POPDC1 expression is not required for TREK-1 association with Ca^2+^/calmodulin-stimulated AC activity

The HR variability displayed upon *Popdc1* deletion is stress-induced, occurring immediately in response to ISO stimulation (*13*) and is a substantially stronger phenotype than that observed upon deletion of AC9. Therefore, we hypothesized that POPDC1 may interact with multiple AC isoforms, accounting for the remaining TREK-1-associated AC activity in *Adcy9*^-/-^ mice (Fig 6A). The calcium-activated AC isoforms, AC1 and AC8, present in atria and SA node, regulate pacemaker activity in the SA node (*28*), and thus represented good candidates to potentially regulate TREK-1 channels. To determine if AC1 or AC8 interacts with POPDC1, non-tagged AC1 and AC8 were expressed with MYC-tagged POPDC1 in HEK293 cells and an IP-AC assay was performed. POPDC1 was weakly associated with AC1 and AC8 Ca^2+^/calmodulin-stimulated AC activity (1.59 +/- 0.05 and 1.8 +/- 0.2 fold over IgG controls, respectively; Fig 6D), demonstrating that multiple cardiac AC isoforms can potentially complex with POPDC1. In cardiac tissue, TREK-1 also pulled down Ca^2+^/calmodulin-stimulated AC activity (2.8 +/- 1.3 fold over IgG controls), suggesting that AC1 and/or AC8 may participate in the regulation of TREK-1 (Fig 6E). However, deletion of POPDC1 did not alter the amount of Ca^2+^/calmodulin-stimulated AC activity associated with TREK-1 in the heart (Fig 6C). Therefore, POPDC1 is required for the association of Gαs-stimulated, but not Ca^2+^/calmodulin-stimulated, AC activity with TREK-1.

## DISCUSSION

AC9 represents the most distantly related of the mammalian AC isoforms but with clear roles in cardiac function (*2*). We show that AC9 is important for resting HR control in mice. Furthermore, we identify POPDC1 as a novel scaffold that mediates interactions of AC9 with the two-pore potassium channel TREK-1 in an ISO-dependent manner. POPDC1 and TREK-1 were previously shown to be important players in HR control, while deletion of *Popdc1, Kcnk2* or *Adcy9* gives rise to HR variability. Our data suggest a model whereby POPDC1 recruits AC9 to a complex with TREK-1 to enhance local cAMP within the complex and facilitate POPDC1 regulation of TREK-1.

### POPDC1 is a scaffold for AC9 complexes

In an effort to examine the molecular mechanism underlying AC9-dependent HR regulation, we identified POPDC1 as a novel AC scaffolding protein that also serves as a cAMP effector involved in HR control. AC9 and AC3 were originally identified with POPDC2 as bait in a large-scale human interactome screen (*30*), but this putative interaction has never been biochemically verified nor subjected to further analysis. We now demonstrate an interaction between AC9 and all three POPDC isoforms using cellular and biochemical methods, including PLA, BiFC, FLIM-FRET, and co-immunoprecipitation followed by western blot and AC activity assays.

POPDC proteins are localized in multiple membrane compartments and have been identified in adult cardiomyocytes in different sarcolemmal compartments including, intercalated disk, t-tubules, caveolae, costameres and the lateral plasma membrane (*35*). POPDC1 can homodimerize, involving residues within the Popeye domain and likely the transmembrane domain (*36, 37*). Given that deletion or mutation of POPDC1 often leads to loss of POPDC2 membrane localization, and deletion of either POPDC isoform closely phenocopies one another with respect to HR variability and stress-induced bradycardia, POPDC1 likely heterodimerizes with POPDC2 (*15, 18, 38*). To avoid possible interference by expression of other POPDC isoforms, AC9 interactions with individual POPDC isoforms by BiFC were performed in COS-7 cells, lacking detectable levels of POPDC1 or −2. However, it is unknown if AC9 interacts only with homodimers or also with potentially existing heterodimeric POPDC complexes.

We used immunoprecipitation of POPDC proteins followed by measurement of the associated AC activity (IP-AC assays) to characterize POPDC-AC interactions in HEK293 cells. Although IP-AC assays have been used to identify numerous AC:AKAP interactions in cells and tissues, an inactive AC was never previously pulled down with an AKAP or PKA effector protein (*10, 39-41*). We show clear interaction of POPDC2 with AC9 by co-immunoprecipitation-western blotting, but were surprised that no Gαs-stimulated AC activity was detected in these pull-downs. POPDC2 may play a second, inhibitory role for AC9. Alternatively, given the possible existence of heteromeric complexes consisting of POPDC1 and POPDC2, fully active AC9 may bind to POPDC1 homodimers and possibly a POPDC1/2 heteromeric complex, while an AC9 enzyme incapable of Gαs stimulation is associated with POPDC2. The complex regulation of AC activity by POPDC proteins remains under investigation.

### TREK-1 associates with multiple AC isoforms in both a POPDC-dependent and independent manner in the heart

TREK-1 channels are mechanoactivated K^+^ channels that are highly expressed throughout the human heart, including the SA and AV nodes (*16, 42, 43*). TREK-1 is thought to regulate the resting membrane potential and therefore determines cell excitability, giving rise to a background K^+^ current in SA nodal cells (*42*). TREK-1 channels are modulated by cAMP via two distinct mechanisms (*44*). TREK-1 *gating* is inhibited by PKA phosphorylation of TREK-1, which is facilitated by AKAP79 association (*45*), while TREK-1 channel density on the plasma membrane is enhanced by POPDC1 (*13, 16*). However, upon POPDC1 binding of cAMP, the TREK-1:POPDC1 complex is dissociated, leading to loss of TREK-1 at the membrane and decreased K^+^ current (*13*).

TREK-1 is co-localized with a complex of POPDC1:AC9 and associates with both POPDC1 and AC9 as revealed by co-immunoprecipitation when expressed in HEK293 cells. POPDC1 and TREK-1 association was measured previously by FRET (*13*). However, the measurement of cellular interactions between AC9 and TREK-1 with FRET-based approaches proved more challenging, due to distance constraints since AC9, POPDC1, and TREK-1 all exist as dimers (*36, 46, 47*). AC9:TREK-1 interactions were detected by BiFC, albeit with smaller signals than those for AC9:POPDC or dimeric complexes of TREK-1, POPDC1, or AC9. Activation of endogenous beta-adrenergic receptors by ISO decreased the AC9:TREK-1 and POPDC:TREK-1 interactions in a dose-dependent manner. This was similar to the decrease in FRET measured in HEK293 cells between POPDC and TREK-1 by ISO, which occurs rapidly and is independent of PKA activity (*13*). BiFC signals for POPDC1:AC9 association were largely independent of ISO treatment, suggesting that this core complex stays intact, while TREK-1 at least partially dissociates.

In the heart, abundant Gαs-stimulated AC activity is associated with TREK-1, which is absent in cardiac tissue from *Adcy9*^-/-^ mice, suggesting that AC9 makes up the majority of TREK-associated AC activity. Deletion of POPDC1 abolished any TREK-1 associated Gαs-stimulated AC activity, consistent with a complex that is dependent on POPDC1. Our BiFC data support this conclusion, where interactions between TREK-1 and AC9 are not detectable in cells lacking POPDC1 expression.

The major Ca^2+^/calmodulin-stimulated ACs, AC1 and AC8, are expressed at low levels in the SA node and possibly atria, maintaining high calcium-stimulated cAMP levels, which are essential for cardiac pacemaking (*25-28, 48, 49*). In HEK293 cells, AC1 and AC8 were associated with POPDC1 in pull-down experiments. In heart, TREK-1 associates with a low amount of Ca^2+^/calmodulin-stimulated AC activity (∼20% of Gαs-stimulated activity), however this was *independent* of POPDC1 expression. Multiple groups have demonstrated AC8, but not AC1, binding to AKAP79 (*39, 50, 51*). AKAP79 is also expressed in SA node, as detected by RNA sequencing (*52*). Therefore TREK-1 association with Ca^2+^/calmodulin AC activity is likely mediated by AKAP79 scaffolding of both TREK-1 and AC8.

### AC9, POPDC1 and TREK-1 participate in heart rate control

We previously reported a bradycardia in *Adcy9*^-/-^ mice under anesthesia (*24*). Using telemetry, a reduced HR was revealed at rest during daytime when the animals are naturally less active. This was not due to AC9 regulation of I_Ks_ currents, which are absent in adult WT mice (*11*). *Adcy9*^-/-^ mice displayed an increase in HR variability that correlated with a decreased HR during the recovery period from high stress, as mimicked by beta-adrenergic stimulation upon ISO injection. However, during exercise, or immediately following ISO injection, the increased HR in *Adcy9*^-/-^ mice was unchanged compared with wild-type. This is consistent with the lack of AC9 regulation of global in vivo PKA phosphorylation of targets such as CREB, troponin I, or phospholamban in response to ISO injection (*24*). Similarly, loss of AC9 decreases the PKA-dependent phosphorylation of the heat shock protein 20 at baseline, but not upon ISO injection. Thus, AC9 represents only a small percentage of AC activity in heart and functions largely at baseline or within local scaffolded complexes (*11, 24*).

TREK-1 is also important for resting HR, as a cardiac specific knockout of TREK-1 in mouse exhibits bradycardia with frequent sinus pauses (*42*). TREK-1 downregulation has been further implicated in atrial fibrillation while a heterozygous point mutation of TREK-1 is linked to idiopathic ventricular arrhythmia (*53*). Mutant mice with a deletion of either *Popdc1* or *Popdc2* exhibit severe stress-induced bradycardia, with high HR variability and long sinus pauses (*13*). In patients carrying missense or nonsense mutations of *POPDC1* that affect cAMP binding or impairs membrane trafficking lead to a host of cardiac and skeletal muscle associated phenotypes (*18*). For example, patients carrying the recessive missense mutation *POPDC1 p*.*S201F*, which affects a residue of the phosphate binding cassette and causes a 50% reduction in cAMP binding, results in impaired TREK-1 regulation in response to ISO, and patients develop a second degree AV block and LGMD (*15*). A dominant nonsense mutation in POPDC2, *POPDC2W188X*, deletes part of the cAMP binding domain and similar to *POPDC1* mutations, leads to a loss of TREK-1 modulation when co-expressed in *Xenopus oocytes*, and causes an AV block in affected patients (*16*). It is unlikely that the impaired regulation of TREK-1 alone is sufficient to explain the strong phenotype of the *Popdc1* and *-2* null mutants, particularly when compared to phenotypes of *Kcnk2* or *Adcy9* deletions in mice. Recently, an interaction of POPDC1 and PDE4 isoforms have been described (*54*). A cell-permeable peptide that disrupts the POPDC1:PDE4 complex caused a reduction in cycle length of spontaneous Ca^2+^ transients in mouse SA nodes. This effect of the peptide was only seen at baseline and was blunted after ISO stimulation. It will be important to determine whether PDE4 is part of the same scaffold as AC9 and TREK-1. POPDC proteins interact with several additional cardiac proteins some of which also have been implicated in cardiac pacemaking, including caveolin-3 (*55*). Mutation of *CAV3* or loss of caveolae gives rise to LGMD and cardiac conduction abnormalities, while deletion of *Cav3* in mice leads to bradycardia (*56*). Multiple AC isoforms are found in caveolae, however, it is unknown whether POPDC proteins are involved in the recruitment of AC proteins to caveolae (or vice-versa). Cardiac pacemaking involves two interacting networks of electrogenic proteins, the membrane and Ca^2+^ clocks (*57*); one or several of these proteins could be modulated through interactions with POPDC proteins. For example, the sodium-calcium exchanger (NCX1) was identified as a POPDC2 interacting protein (*58*) and ablation of *Slc8a1* was shown to cause SAN dysfunction (*59*). Whether NCX1 is also part of the POPDC1:AC9:TREK-1 complex needs to be tested. It is also unknown whether AKAP proteins are recruited to POPDC-scaffolded complexes via association with AC. AKAP79 is not detected with TREK-1:POPDC1:AC9, however that does not rule out the possibility of AKAP recruitment to other POPDC:AC-containing complexes. POPDC1 has also been implicated as a negative regulator of hippocampal synaptic plasticity (*60*). Interestingly, both TREK-1 and AC9 are expressed in the hippocampus (*61*), with TREK-1 also regulating synaptic plasticity (*62*) and thus, the AC9:POPDC1:TREK-1 signaling complex may be present in other cell types and in different physiological contexts. In conclusion, POPDC proteins represent a unique cAMP effector and AC9 scaffolding protein that bridge interactions with TREK-1 to regulate heart rate.

## METHODS

### Gene-Targeted Mice

*Adcy9* knockout mice were obtained from the Mutant Mouse Regional Resource Center. The mouse strain B6;129S5-Adcy9Gt(neo)159Lex/Mmucd, identification number 011682-UCD contains an insertion of the gene trap vector between exons 1 and 2 of *Adcy9*; generation, backcrossing and genotyping of mice are further described in (*24*). *Adcy9*^-/-^ mice show a preweaning sub-viable homozygous phenotype with incomplete penetrance. To obtain the necessary animal numbers, crosses were set up with *Adcy9*^-/-^ x *Adcy9*^+/-^ versus WT x *Adcy9*^+/-^. Age matched, wild-type C57BL/6J controls were used for all behavioral experiments. Animals were housed at 22°C with a 12 h light/dark cycle (7AM-7PM) with free access to water and irradiated chow diet. For exercise experiments, animals were not pre-trained to run. All *Adcy9*^-/-^ animal protocols were approved by the Institutional Animal Care and Use Committee (IACUC) at the University of Texas Health Science Center at Houston in accordance with the Animal Welfare Act and NIH guidelines. *Popdc1*^-/-^ mice were generated by replacement of the first coding exon with a β-galactosidase-encoding gene cassette as described (*63*). The *Popdc1*^-/-^ mice were backcrossed and maintained on a C57BL/6J background. Wild-type controls for biochemical assays of hearts from *Adcy9*^-/-^ or *Popdc1*^-/-^ mice used age-matched and/or littermates from their respective breeding facilities. Work with *Popdc1*^-/-^ animals was approved by the Animal Welfare and Ethical Review Board of Imperial College London.

### Behavioral Assays

All tests were performed by blinded investigators. Mice were acclimated to the behavioral testing room for 30–60 minutes in their home cages with the investigator present before beginning testing. Elevated plus maze and open field activity box was performed one week prior and again after telemetry implantation surgery.

### Elevated Plus Maze

The 40 cm high EPM was used to measure differences in anxiety. It consisted of four, 12 cm wide arms: two enclosed by 40 cm high walls and two open (modeled upon the Stoelting Co’s EPM model, Wood Dale, IL). Individual mice were placed in the EPM center facing an open arm and explored freely for 5 minutes. The test was video recorded and later analyzed for time spent in the open versus closed arms by an investigator blinded to genotypes, as previously described (*64-66*).

### Open-Field Activity Box

The open-field box (ENV-515, Med Associates, Inc., St. Albans, VT) consists of an activity chamber (43.2 cm wide by 43.2 cm deep by 30.5 cm high) with 16 infrared transmitters and receivers evenly positioned around the chamber’s periphery. Mice placed in the activity chamber moved freely for 25 minutes. Data were collected by the Open Field Activity (OFA) software that registers time spent and movements within the chamber as a whole versus the inner zone 1 by recording photobeam interruptions.

### ECG and telemetry

ECGs were recorded using an ETA-F10 ECG transmitter (DSI, St. Paul, MN) implanted into 10 wildtype and 10 AC9 KO mice (6 males and 4 females per group; one male WT mouse removed the transmitter lead and was dropped from all telemetry analysis). Mice were allowed to recover for at least one week post-surgery. ECGs were recorded in mice 5-8 months of age (average weight minus probe = 24.4 +/- 2.7 g). Telemetric ECG was recorded during normal activity and during the following stress tests: free running wheel, forced swim test, and ISO injections (methods described below). ECG data was analyzed in Ponemah v6.52 (DSI, St. Paul, MN). HR and RR intervals were averaged from Ponemah exported data. Telemetric ECGs were manually analyzed with respect to arrhythmias by examiners blinded to the genotypes and corrections made for any beats not correctly identified by the automated software analysis. HR variability was calculated in Ponemah using the Variability Analysis Time Domain function and averages were determined from the exported data from Ponemah. pNNx value was set at 6 ms, as is standard for mice.

### Running Wheel

Mice were placed in cages for 24 hrs containing voluntary running wheels fitted with electronic monitors for activity tracking by Activity Wheel Monitor software (Lafayette Instruments). ECGs were recorded throughout the 24 hr period. Distance traveled on wheels was cross referenced with the HR data to calculate HR during active versus inactive and recovery periods. Active times were defined as sustained activity on the wheel for 5 minutes and selected by examiners blinded to the genotypes. Inactive times were selected only if the wheel did not move for a full 10 minutes. Active and inactive times were selected for analysis by blinded investigators.

### Forced Swim Test

Forced swim tests were conducted in 5 liter beakers filled with autoclaved water (26-28 degrees C) to 16 cm according to published protocols (*67*). Mice were placed in water containers for 6 minutes; recorded swimming behavior was quantified during the last 4 minutes by blinded investigators.

### ISO Injections

ECGs were recorded for at least one hour prior to injections. ISO (1 µg drug per g weight of mouse; subtracting probe weight) was administered by intraperitoneal injection. Mice were returned to cages for at least 1 hour and 30 minutes post ISO injection for ECG recordings.

### Plasmids

Flag- and YFP-tagged AC9 pCDNA3 plasmids are as described (*24*). POPDC1-myc, POPDC2-myc and POPDC1Δ172-myc were described in (*13, 15*). Bimolecular Fluorescence complementation constructs consist of split Venus fluorescent protein. The N-terminal (VN) or C-terminal (VC) half of Venus was fused to the C-terminus of mouse POPDC1, POPDC2, POPDC3, and POPDC1Δ236, and the N-terminus of human TREK-1c. In each case, previously described tagged constructs of each protein were used as the starting point for cloning, whereby the existing tag was swapped for a BiFC tag by PCR (*13, 15*). AC9-VN and AC9-VC lack N-terminal tags and contain a 7 aa linker (AAAGGGS) between AC9 and VN or VC, as described (*47*).To create the myristoylated cytoplasmic domain of POPDC1, the myristoylation and palmitoylation sequence from Lyn was fused to aa 117 of POPDC1 (starting immediately after the three membrane-spanning regions); a Myc- or VN-tag is contained at the C-terminus. All constructs were verified by DNA sequencing.

### Cell culture and transfections

HEK293 and COS-7 cells were authenticated by ATCC, cultured in Dulbecco’s Modified Eagle Medium with 10% fetal bovine serum, and transfected with the indicated plasmids using Lipofectamine 2000 (*39, 41*). ISO was stored and diluted in AT buffer (100 mM ascorbate and 10 mM thiourea, pH 7.4) for cellular studies.

### Proximity Ligation Assay

In situ PLA was performed using a Duolink kit (Sigma-Aldrich, cat. DUO92101) following the manufacturer’s protocol. HEK293 cells were cultured on clear bottom 96 well plates (Greiner Bio-One), transfected with the required plasmids and fixed with 4% PFA. After washing the plate 3 times with PBS, the cells were blocked (1% BSA + 0.075% Triton X100) for 1 h at room temperature and then incubated with primary antibodies overnight. Antibodies included: rabbit anti-GFP (SC-8334, SantaCruz, 1:1000) and mouse anti-MYC (purified by National Cell Culture from the ATCC hybridoma CRL-1729 for MYC 1-9E10.2, 1:1000) or mouse anti-Gβγ (SC-378, SantaCruz, 1:1000). After removal of primary antibodies, the samples were incubated with anti-mouse PLUS and anti-rabbit MINUS PLA probes for 1 h at 37 °C. Subsequent steps of ligation and amplification were according to the manufacturer’s protocol. After the last wash, cells were stained with DAPI (1 μg/ml) and imaged using an epifluorescence high content imaging microscope with a 20X objective, 25 fields of view per condition (>2000 cells per condition per experiment; CellInsight CX5 High Content Screening platform, ThermoFisher). Data analysis of mean fluorescence intensity per cell was performed using FACS analysis software (FlowJo, USA). Each experimental condition was repeated at a minimum in 3 separate experiments. To prevent false positives, cells with saturating YFP fluorescence were not considered in the analysis.

### Bimolecular Fluorescence Complementation (BiFC)

Cells were plated in uncoated 12-well plates one day prior to transfection with Lipofectamine 2000. 48 h post transfection, cells were washed once with PBS and then incubated with Tyrode’s buffer at 37 °C for 10 min. Where indicated, cells were treated with or without ISO at 37 °C for an additional 10 min. Cells were harvested and transferred to a 96-well plate; an aliquot of each condition was used for total protein determination by Bradford analysis. BiFC (Venus) signals were obtained using a Tecan fluorescence microplate reader at excitation wavelength 506 nm and emission wavelengths 538-542 nm as described (*47*). Background fluorescence was averaged from multiple wells with cells expressing pCDNA3 and proteins tagged with VN or VC in the absence of the other Venus half. BiFC was calculated as YFP signals minus average background signal and normalized to total protein for each sample in HEK293 cells. For COS-7 cells, 48 h post transfection, cells were washed and incubated with DMEM containing DAPI (10 µg/ml) for one hour at 37°C. Cells were washed, transferred in PBS to 96-well plates, and YFP and DAPI (358 nm excitation and 461 nm emission) signals were measured at room temperature. YFP signals were normalized to DAPI staining after subtraction of average background YFP fluorescence from multiple control samples.

### Live-Cell Imaging

For BiFC imaging, HEK293 cells were plated on Matek dishes coated with poly-lysine. Prior to imaging, media was replaced with Tyrode’s buffer (10 mM HEPES, pH 7.4, 145 mM NaCl, 4 mM KCl, 1 mM MgCl2, 1 mM CaCl2, 10 mM D-Glucose). BiFC images were acquired using a TE 2000 microscope (Nikon, Tokyo, Japan) with a DG4 xenon light source and CoolSNAP cameras (Roper Scientific, Trenton, NJ). Venus fluorescent protein images were acquired 36-48 hours after transfection (excitation 500/20 nm, emission 535/30 nm).

### FLIM-FRET

FLIM experiments were performed with a lifetime fluorescence imaging attachment (Lambert Instruments, Leutingewolde, the Netherlands) on an inverted wide-field microscope (Nikon). Cells plated on poly-lysine coated Matek dishes and transfected as indicated were excited by using a sinusoidally modulated 3-W 448-nm light-emitting diode at 40 MHz under epi-illumination. Fluorescein at the concentration of 1 µM was used as a lifetime reference standard (4 ns). Cells were imaged with a 40x Plan-Apo oil-immersion objective (numerical aperture 1.45) using a CFP filter set; exposure times were less than 150 ms to avoid photobleaching. The phase and modulation lifetimes were determined from a set of 12 phase settings using the manufacturer’s software (LIFA). FRET efficiency was calculated according to formula F_eff_ = 1 - τ2/τ1. FLIM data were averaged on a per-cell basis. Three experiments (>40 cells in total) were performed for each condition. Donor alone and donor-acceptor measurements were taken immediately after one another. In order to prevent differences in donor lifetimes due to altered lipid environments, Cerulean-TREK-1 lifetime was measured in the presence of POPDC1 (tagged with Myc or Venus) to enhance PM localization.

### Immunoprecipitations and Western blotting

Antibodies and reagents used for immunoprecipitation and western blotting include mouse anti-FLAG M2 agarose affinity gel (Sigma-Aldrich), mouse anti-DYKDDDDK (Flag) tag (Cell Signaling Technologies, Danvers, MA), mouse anti-MYC (purified by National Cell Culture from the ATCC hybridoma CRL-1729 for MYC 1-9E10.2), mouse anti-A.v. monoclonal antibody for green fluorescent protein (JL-8; Takara Bio, Kusatsu, Japan; recognizes VC), rabbit anti-GFP (D5.1 Cell Signaling Technology 2956S; recognizes VN), mouse anti-β-actin (C4, Santa Cruz Biotechnology), anti-POPDC1 (early studies used Santa Cruz, goat #sc-49889 but now discontinued; Sigma-Aldrich, rabbit #HP018176), and normal mouse or rabbit IgG (Santa Cruz Biotechnology). The rabbit anti-AC9 antibody was generated and characterized as described (*47*).

### Adenylyl Cyclase Activity and Immunoprecipitation-AC Assays

Preparation of heart extracts and measurement of AC activity were performed as previously described (*39, 41*). Immunoprecipitation of POPDC complexes followed by western blotting or measurement of associated AC activity (IP-AC assay) was performed as described (*68*). AC activity was stimulated with the indicated reagents or proteins (purified and activated GTPγS-Gαs or calcium and purified calmodulin) and cAMP was detected by enzyme immunoassay (Assay Designs) or using [γ^32^P]ATP.

### Statistical Analysis

Repeated Measures ANOVA was used for testing the significance in the HR and HR variability data with multiple data points over time. Data are expressed as mean ± standard error of the mean (SEM), except for behavioral data in Table I which are expressed as mean ± the standard deviation (SD). Normality was determined by Shapiro-Wilk test. Differences between samples were determined using one-way or two-way analysis of variance (ANOVA) followed by the indicated tests for comparison between multiple groups. For comparisons between two groups with equal variance, Student’s t test was used. For samples of non-equal variance, the non-parametric Kruskal-Wallis ANOVA on Ranks or Mann-Whitney Rank Sum test was performed. Significant p values are indicated as follows: (*) denotes a p value < 0.05, (**) < 0.01 and (***) < 0.001. All analyses were performed using SigmaPlot statistical analysis software.

## Acknowledgements

We wish to thank Simran Rahman and Annabel King for their blinded analysis of behavioral videos and telemetry data and Ursula Herbort-Brand for her excellent technical assistance.

## Funding

National Institutes of Health grant RO1GM60419 (CWD)

British Heart Foundation grants PG14/46/3091 and PG19/13/34247 (TB)

National Institutes of Health grant R37NS096493 - use of telemetry equipment

## Author contributions

Designed the research: TAB, YL, TB, and CWD

Performed the research: TAB, YL, AM, MZ, AGC, and MAG.

Performed surgeries for telemetry implantation: VRV

Analyzed the data: TAB, YL, AM, AGC, and CWD

Provided key reagents: TB and RFRS

Supervised the study: TB and CWD

Wrote the manuscript with edits by all authors: YL, AM, and CWD

## Competing interests

The authors declare that they have no competing interests.

## Notes

### Competing Interest Statement

The authors have declared no competing interest.

